# Mental Rotation is a weak measure of the propensity to visualise

**DOI:** 10.1101/2024.10.29.620977

**Authors:** Derek H. Arnold, Loren N. Bouyer, Blake W. Saurels, Elizabeth Pellicano, D. Samuel Schwarzkopf

**Affiliations:** Perception Lab, School of Psychology, The University of Queensland; Clinical, Educational and Health Psychology, University College London; Experimental Psychology, University College London; School of Optometry & Vision Science, The University of Auckland

**Keywords:** Visual Imagery, Mental Rotation, Aphantasia

## Abstract

There is increasing evidence of substantial differences in people’s capacity to voluntarily visualise – with some (Congenital Aphants) asserting they cannot visualise at all. Its been suggested that Congenital Aphants might be mistaken about their inability, as some have performed similarly on tasks purported to measure imagery – including the mental rotation task, where people decide if objects depicted from different viewpoints are the same or different. We examined how the vividness of people’s imagery is related to performance on a ‘mental rotation’ task. People also reported on their response strategies. Mental rotation was an overall superior response strategy relative to non-visualising. However, the vividness of people’s imagery was only weakly associated with viewpoint contingent changes in task performance, and it did not predict changes in reliance on mental rotation as a response strategy. Overall, our data suggest performance on mental rotation tasks is a weak measure of people’s propensity to visualise.

---

Aphantasia is a term coined to refer to an inability to have voluntary imagined visual sensations – or to visualise (Zeman et al., 2010; Zeman et al., 2015). There is good evidence that Aphantasia is associated with behavioural (e.g. Chang & Pearson, 2018; Keogh & Pearson, 2011, 2014; Jacobs et al., 2018; Liu & Bartolomeo, 2023; Milton et al., 2021), physiological (e.g. Kay et al., 2022; Wicken et al., 2021) and with morphological differences to the structure of human brains (Bergmann et al., 2016; Milton et al., 2021).

Some researchers have suggested that Aphants might be mistaken about the nature of their inability (e.g. Nanay, 2021; Siena & Simons, 2024; Pounder et al., 2022). This conjecture has been motivated by findings that some Aphants have performed similarly to non-aphants on tasks that purportedly measure the propensity to visualise (e.g. Jacobs et al., 2018; Pounder et al., 2022; Zhao et al., 2022). In a mental rotation task, for instance, people can be asked to decide if adjacent images depict the same or different (often mirror reversed) objects shown from the same or from different viewpoints (Shepard & Metzler, 1971; Ganis & Kievit, 2015; see Figure 1). Many people report using a strategy where they visualise one of the two objects rotating in their mind’s eye, until it is seen from the same perspective as the other image – thereby helping them to decide if the two images depict the same or different objects. Researchers have regarded this task as a metric of people’s capacity to visualise (e.g. Just & Carpenter, 1985; Shepard & Metzler, 1971). Consequently, findings that Aphants can perform similarly on mental rotation tasks (e.g. Jacobs et al., 2018; Marks, 1999; Pounder et al., 2022) have caused researchers to question the nature of aphantasia – with some suggesting that Aphants might be able to visualise but lack awareness of this capacity (Nanay, 2021; Siena & Simons, 2024; Pounder et al., 2022). We thought another possibility should be considered. Perhaps mental rotation tasks are an unreliable metric of people’s propensity to visualise (also see Marks, 1999)?

**Figure 1.**
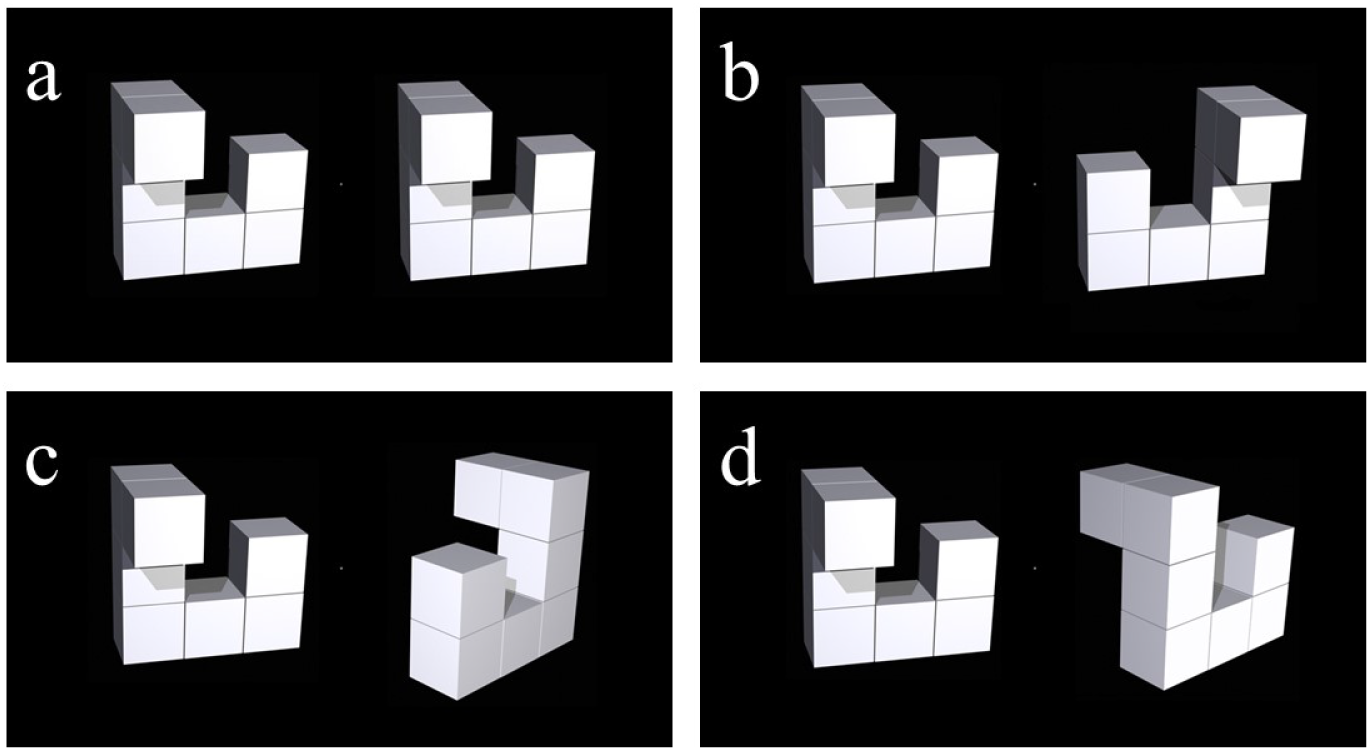
**a)** Images depicting the same object viewed from the same viewpoint. **b)** Images depicting mirror-reversed objects, viewed from the same viewpoint. **c)** Images depicting the same object, viewed from viewpoints rotated 150°. **d)** Images depicting mirror-reversed objects, viewed from viewpoints rotated 150°. All images are taken from a set of images depicting three-dimensional shapes for investigating mental rotation – published by Ganis & Kievit (2010).

When people have been quizzed about how they perform mental rotation tasks, a number of strategies have been identified – including analytic response strategies that do not involve visualisation (Bethell-Fox & Shepard, 1988; Hegarty, 2018; Khooshabeh et al., 2013).

Indeed, evidence suggests that, in some circumstances, a non-visualising strategy can result in *superior* performance on a ‘mental rotation’ task (Hegarty, 2010). A very recent study compared the performance of a group of Aphants on mental rotation tasks to a control group who could visualise. They found that Aphants were overall slower, but no less accurate in terms of their task performance (Kay et al., 2024). Participants were also quizzed about their response strategies, and Aphants more often reported using non-visualising analytic strategies. As the difference in decision speeds could be attributed to Aphants having taken more caution, there is some ambiguity about how to interpret this evidence – did the Aphants perform the task in a qualitatively different way, or were they just more cautious? Regardless of this interpretive ambiguity, the authors highlighted the differences in subjective response strategies, and argued that their data show that similar ‘mental rotation’ performance levels can be achieved using different response strategies. Findings like this seem sufficient to call into question the utility of mental rotation tasks as a reliable metric of people’s propensity to visualise (also see Khooshabeh et al., 2013), particularly if people are not asked about their response strategies.

Here we ponder the following question. Are mental rotation tasks generally a reliable estimate people’s propensity to visualise? We had people perform a mental rotation task, and report on their response strategies. To preface our results, while we find that people who report having vivid imagery are more likely to report visualising an object rotating, and they make proportionally fewer incorrect decisions about objects depicted from different viewpoints, these associations are weak. Moreover, people often report both visualising when this is disadvantageous (when mirror-reversed objects are depicted from the same viewpoint), and that they have relied on non-visualising analytic response strategies.

## Materials and Methods

All data relating to this study will be made available via UQ eSpace.

### Ethics

Ethical approval was obtained from the University of Queensland’s (UQ) Ethics Committee. The experiment was performed in accordance with UQ guidelines and regulations for research involving human participants. Each participant provided informed consent to participate, and they were informed that they could withdraw from the study at any time without prejudice or penalty.

### Participants

A total of 231 people commenced, but 40 failed to complete the study. We excluded a further 96 people on the following basis. The task response sequence was 2-part, with an additional response required when people reported that they had ***not*** visualised a rotation. This meant that there was an incentive to report visualisations to finish the study quickly. We therefore excluded *all* participants who had either failed to complete the study, or who had reported visualising an object rotating when test images had depicted the same object seen from the same viewpoint.

The final sample size included 95 participants (M age = 20, SD = 4.2, range = 17 – 38). Of these participants, 67 identified as women, 27 as men, with 1 ‘other’. All participants were undergraduate students at The University of Queensland, who took part in the study for course credit.

### Stimuli

This study was conducted online, via Qualtrics (Qualtrics, Provo, UT). After providing informed consent to take part, participants were first asked to respond to some demographic questions (Age, Gender identity). After, they completed Mental Rotation trials, followed by the Vividness of Visual Imagery Questionnaire v2 (VVIQ-2, Marks, 1995), which provides a subjective assessment of the vividness of a participant’s imagined visual experiences.

Images for the mental rotation task were selected from an image set depicting three-dimensional shapes (Ganis & Kievit, 2015). There were four conditions, with pairs of images depicting the same or different (mirror reversed about the horizontal axis) objects, depicted from the same or from different viewpoints (i.e. rotated horizontally by 150°) – a 2 x 2 design (see Figure 1 for an example of each type of test image). We selected 8 images for each experimental condition, so there were 32 trials in total, which were all completed in a randomised order.

### Procedure

Before mental rotation trials, participants were given the following instructions:

> “You are about to see a number of pictures. Each depicts two objects. You task is to decide if they are the same object, depicted twice, or if the two objects are different.

> *Sometimes the two objects will be shown from the same angle, and sometimes the object on the right will be rotated relative to the object on the left. When the two objects are shown from different angles, half the time they will be the same object, and half the time they will be different objects*.

> *Many people find it helps to imagine one of the two objects rotating in their mind, to see if it matches the other object after they have seen it rotate. Other people rely on an intuition, that the objects are the same or different. Still other people judge if prominent features and turning points of the objects would match if one of the two objects were physically turned - but they do not visualise either object as having turned*.

> *Your task on each trial is to judge as quickly and as accurately as possible whether the two objects are the same or different. You will then be asked how you came to that decision.”*

After reading these instructions, participants completed 4 practise trials, one for each condition. On practise trials feedback was given regarding the accuracy of same / different decisions. Participants received no feedback on experimental trials. The images used for practise trials were not re-used in the formal experiment.

Test image presentations were shown until the participant had chosen the ‘same’ or the ‘different’ response option, and had then clicked on an advance icon. The timing of the last response option selection (participants could change their responses) was taken as the response time for that trial (not the timing of the advance icon click). After this response sequence, participants were asked “*Did you visualise (have an imagined visual experience of) one of the objects rotating while making your decision?*”. If they responded *No*, they were additionally asked “*Which of these best describes how you made your decision”*. There were four response options for this question, **1)** *I could immediately tell*, **2)** *I guessed*, **3)** *I made a careful cross comparison of the two objects’ features (without having any imagined experience)*, and **4)** *I used a strategy not described here*. If participants answered 4), at the end of the block of trials they were reminded that they had reported using a strategy that we had not described, and they were asked to describe that strategy (or strategies) in a text entry box.

## Results

In all correlational analyses, we excluded data points based on Mahalanobis distance calculations, using a significance level of 0.9. We also conducted a Shapiro-Wilk’s test to assess if each dataset conformed with the normality assumption. When they did, a Pearson’s correlation coefficient was calculated; otherwise, a Spearman’s rank correlation coefficient was calculated. Analyses of response times (RTs) related to individual conditional median RTs for correct decisions.

### People were slower and less accurate when making decisions about rotated objects

First, we address commonly accepted hallmarks of mental rotation – the increased time and difficulty associated with evaluating objects depicted from different, as opposed to the same viewpoint. Consistent with these, we found that individual median response times (RTs) were *longer* when objects were depicted from different viewpoints (M = 4.86, SD = 2.13) compared to objects depicted from the same viewpoint (M = 2.22, SD = 0.8, paired t_94_ = 15.5, p < 0.001). Similarly, when objects were depicted from different viewpoints, people made fewer correct same / different object decisions (M = 0.86, SD = 0.15), compared to when objects were depicted from the same viewpoint (M = 0.97, SD = 0.06, paired t_94_ = −9, p < 0.001).

### Participants did not always report visualising objects rotating, even on harder trials

Next, we highlight that considerable individual differences in the proportions of trials on which visualisations of object rotation were reported. The average proportion was 0.54 (SD = 0.17) overall (see Figure 2a), and 0.83 (SD = 0.24) on trials when objects had been depicted from different viewpoints (horizontally rotated by 150°, see Figure 1c-d and 2b). Participants reported more visualisations of rotation when objects were depicted from different as opposed to the same viewpoint (paired t_94_ = 0.29, p < 0.001).

**Figure 2.**
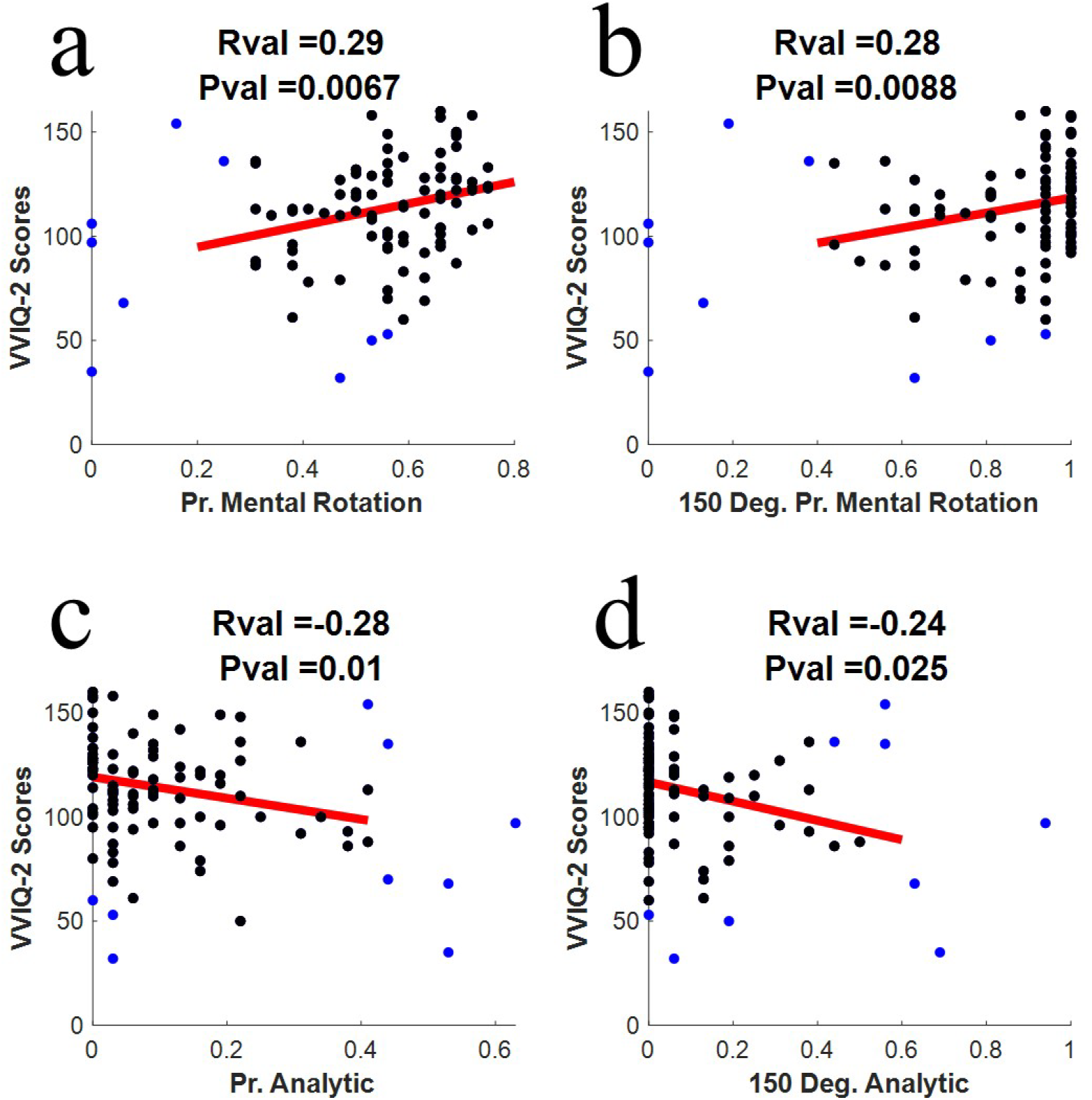
**a)** X/Y scatter plot of the proportion of VVIQ-2 scores (Y-axis) expressed as a function of the proportion of all trials on which participants reported having visualised a test image rotating (X-axis). **(b)** Details are as for Figure 2a, except that data on the X axis depict the proportion of 150° trials on which people reported visualising an object rotating. **(c-d)** Details are as for Figure 2a-b, except that data on Y axes relate to proportions of trials on which people reported using an analytic response strategy. In each plot, datapoints excluded from analyses based on Mahalanobis distance calculations are coloured blue. Red lines depict a linear least squares fit to black data points.

The majority of participants (73%) reported using a non-visualising analytic response strategy on a subset of trials. We estimate people’s propensity to use an analytic response strategy from the proportion of trials on which they endorsed the response option ‘*I made a careful cross comparison of the two objects’ features (without having any imagined experience)*’. Overall, people reported using an analytic response strategy on 0.12 of trials (SD = 0.14; see Figure 2c), and on 0.10 (SD = 0.18) of trials where objects had been shown from different viewpoints (see Figure 2d). There was no robust difference between the proportions of trials on which participants reported using an analytic response strategy when objects were depicted from different as opposed to the same viewpoint (paired t_94_ = 1.57, p = 0.121).

### VVIQ-2 scores and response strategies

Overall, people were weakly *more* likely to report having visualised an object rotating if they had also reported experiencing vivid imagery (r = 0.29, p = 0.007, see Figure 2a). People were also weakly more likely to report visualising an object rotating when objects were depicted from different viewpoints (r = 0.28, p = 0.009, see Figure 2b).

People who had reported having vivid imagery were also weakly *less* likely to report using an analytic response strategy if they (r = −0.24, p = 0.025, see Figure 2c) and they were weakly less likely to report using an analytic response strategy when making decisions about objects depicted from different viewpoints (r = −0.24, p = 0.028, see Figure 2d). Note that these data are not simply the inverse of people reporting on visualising an object rotating as there were other response strategies (i.e. people could report that they had immediately known, or that they had guessed).

### VVIQ-2 scores and task performance

In terms of the speed of responses, there was *no* robust evidence that people who report having vivid imagery are advantaged or disadvantaged when deciding if objects depicted from different, as opposed to the same viewpoint, are the same or different (r = 0.13, p = 0.19; see Figure 3a). In terms of the accuracy of decisions, there was a *weak but robust* relationship between the vividness of people’s imagery (as indexed by VVIQ2 scores) and their accuracy when deciding if objects depicted from different viewpoints were the same or different (r = 0.3, p = 0.003; see Figure 3b).

**Figure 3.**
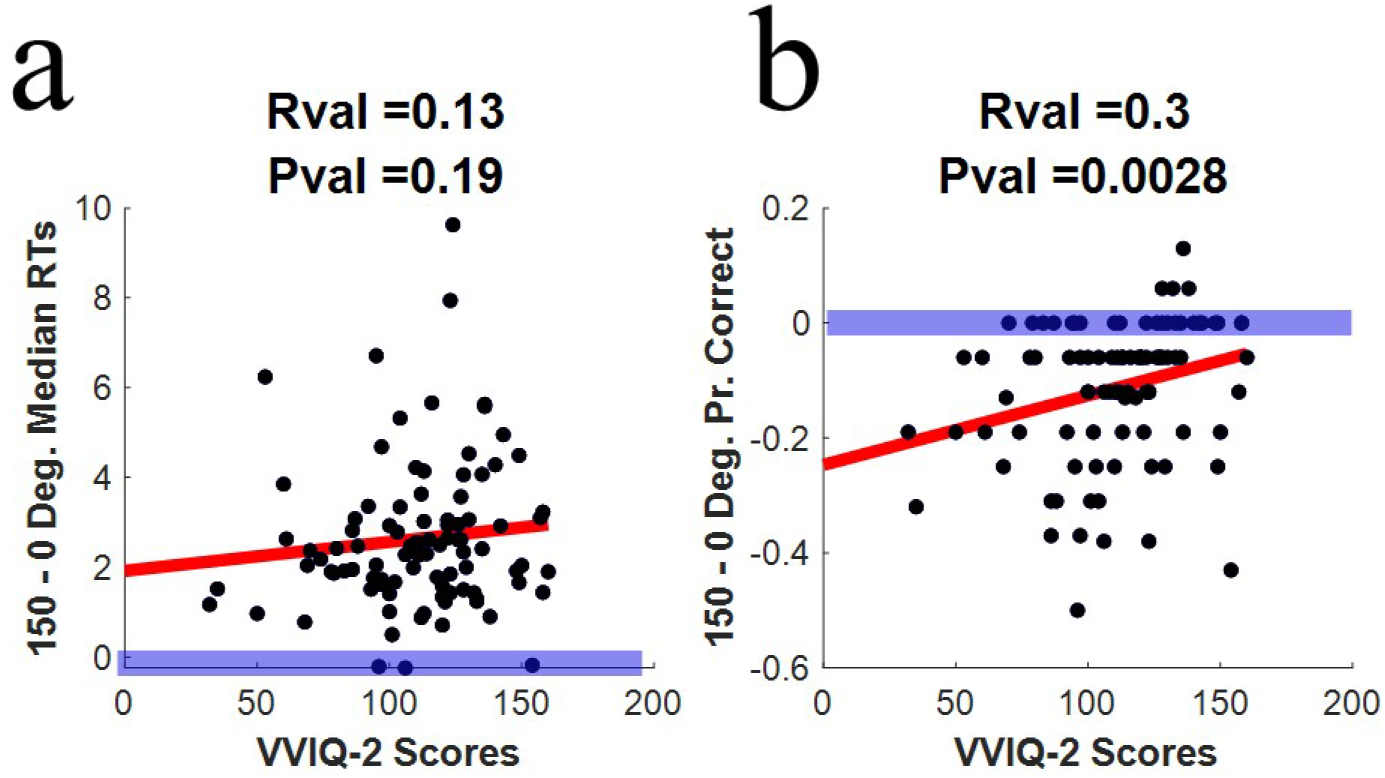
Details are as for Figure 2, with the following exceptions. **(a)** X/Y scatter plot of differences in median RTs when objects were depicted from different (150° horizontally rotated) as opposed to the same viewpoint (Y-axis) and VVIQ-2 scores (X-axis). **(b)** X/Y scatter plot of differences in proportions of correct same / different object decisions when objects were depicted from different (150° horizontally rotated) as opposed to the same viewpoint (Y-axis) and VVIQ-2 scores (X-axis). Bold horizontal blue lines depict no viewpoint differences, either in terms of RTs or in the proportion of correct decisions.

### Changes in response strategy and changes in task performance

Here, we ask if viewpoint dependent changes in response strategy were associated with changes in task performance. Note that these analyses do not reference the typical vividness of people’s imagery (VVIQ2 scores).

There was no robust evidence for a relationship between viewpoint dependent changes in the likelihood of people reporting mental rotation and RTs (r = 0.034, p = 0.76; see Figure 4a). There was, however, a robust *positive* relationship (r = 0.31, p = 0.004; see Figure 4b) between an increased likelihood of people reporting mental rotation and people making fewer incorrect decisions, when objects were depicted from different as opposed to the same viewpoint. This evidence suggests that mental rotation is a superior response strategy to alternatives when people must decide if objects depicted from different, as opposed to the same viewpoint, are the same or different.

**Figure 4.**
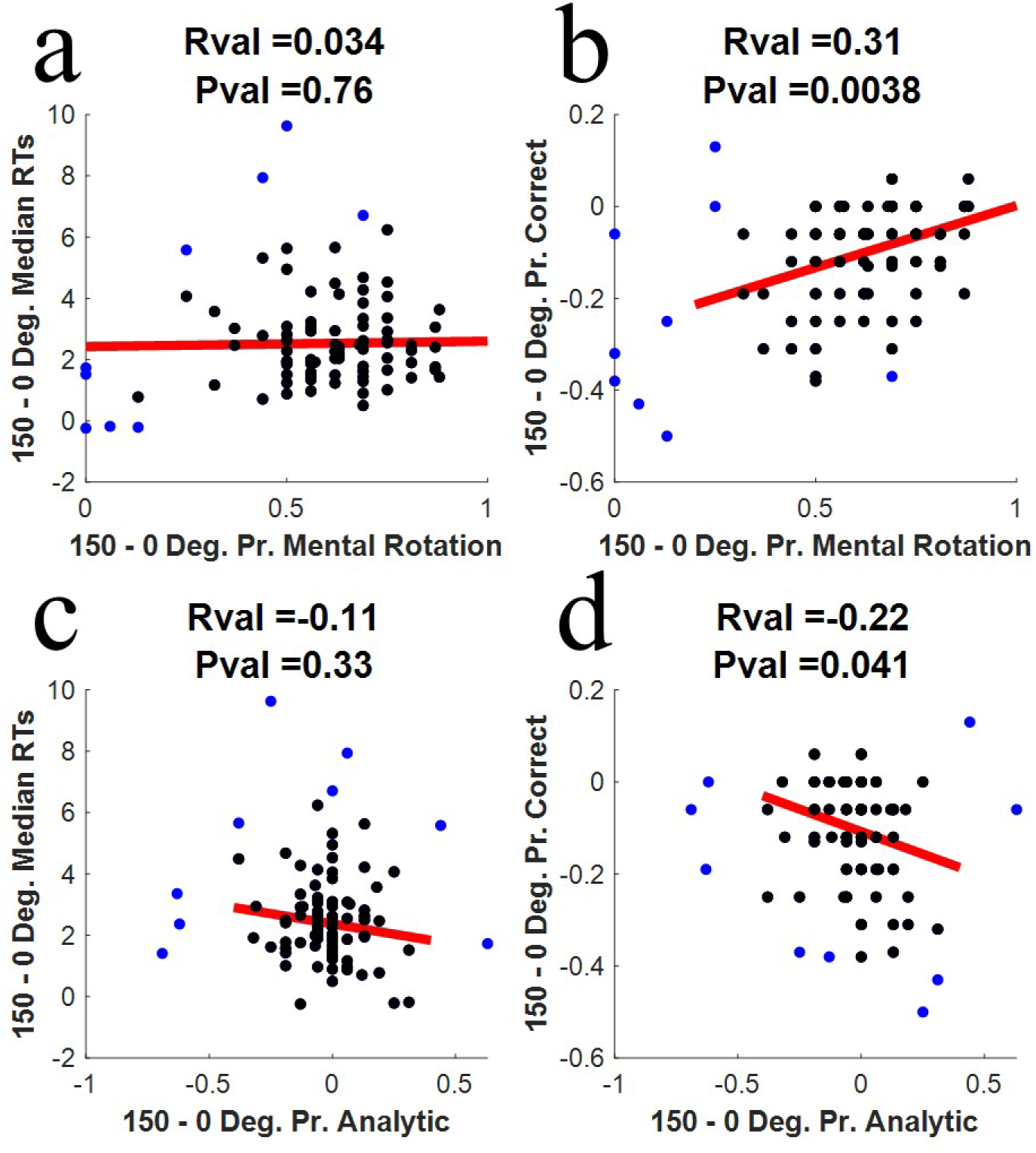
Details are as for Figure 2, with the following exceptions. **(a)** X/Y scatter plot of viewpoint dependent changes in reported proportions of mental rotations (X-axis), and viewpoint dependent changes in median RTs (Y-axis). **(b)** Details are as for Figure 4a, except that data on the Y axis depicts viewpoint dependent changes in correct same / different object decisions. **(c-d)** Details are as for Figure 4a-b, except that data on X axes relate to viewpoint dependent changes in the proportions of trials on which people reported using a non-visualising analytic response strategy.

There was no evidence of a robust relationship between viewpoint dependent changes in reliance on a non-visualising analytic response strategy and RT changes (r = −0.11, p = 0.33, see Figure 4c). There was, however, a *robust negative* relationship (r = 0.22, p = 0.041; see Figure 4d) between an increased likelihood of people reporting that they had relied on a non-visualising analytic response strategy and viewpoint contingent changes in decision accuracy. This suggests that non-visualising analytic response strategies are inferior to alternatives when people must decide if objects depicted from different, as opposed to the same viewpoint, are the same or different.

### Response times and response strategies

Our analyses suggest that viewpoint dependent changes to decision RTs are unrelated to either the vividness of people’s imagery (see Figure 3a) or to changes in their response strategies (see Figure 4a and 4c). One plausible explanation is that these different response strategies have not resulted in different RTs. To address this possibility, we selected all participants available for data analysis who had reported using both strategies for each viewpoint condition, and compared their RTs across the different strategies via a Bayesian paired t-test. This analysis showed that RTs associated with decisions involving mental rotation were interchangeable with RTs associated with decisions involving a non-visualising analytic response strategy, both when objects were depicted from the same viewpoint (t_52_ = 1.528, p = 0.133, BF_10_ = 0.348, see Figure 5a), and when objects were depicted from different viewpoints (t_52_ = 0.321, p = 0.751, BF_10_ = 0.119, see Figure 5b). So, while reliance on a non-visualising response strategy predicts *worse* outcomes in terms of accuracy (see Figure 4d), these inaccurate decisions did not take longer to arrive at.

**Figure 5.**
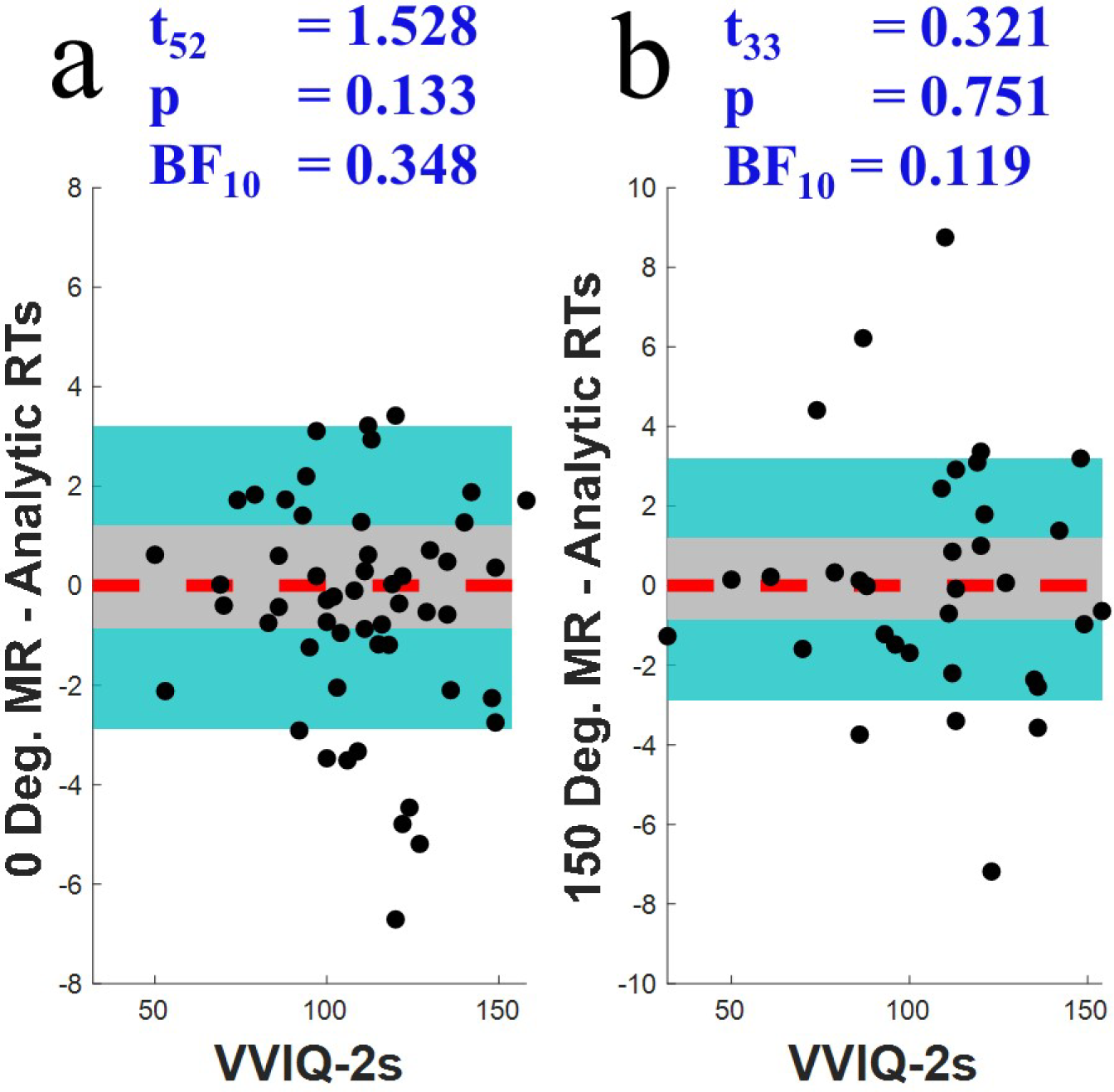
**a)** X/Y scatter plot of differences in individual median RTs, when participants have reported visualising an object rotating as opposed to when they have reported relying on a non-visualising analytic response strategy (Y-axis). Data points are shown in relation to VVIQ-2 scores (X-axis) primarily because this was a convenient way to scatter the individual data points. The red dotted horizontal line depicts no response-strategy dependent differences in RTs. The grey shaded region depicts +/- 2 S.E.s These data relate to trials with objects depicted from the same viewpoint. **(b)** Details are as for Figure 5a, but these data relate to trials where objects were depicted from different viewpoints.

### Viewpoint dependent changes to response strategy are unrelated to VVIQ-2 scores

Our data suggest that any viewpoint related increase in the likelihood of mental rotation *decreases* the number of incorrect same / different object decisions when objects are depicted from different, as opposed to the same viewpoint (see Figure 4b). Our data also suggest that people who report typically experiencing more vivid imagery are more likely to report visualising an object rotating, both overall (see Figure 2a) and when objects are depicted from different viewpoints (see Figure 2b). From this, one might suspect that people who report experiencing more vivid imagery would show more evidence of a strategy shift between viewpoint conditions. Our data speak against this.

In Figure 6a, we depict data describing viewpoint dependent changes in the balance of response strategies. For these data we calculate difference scores relating to the probability that people will report visualising when objects are depicted from different, as opposed to the same viewpoint. We then calculate difference scores relating to the probability that people will report a non-visualising analytic response strategy, and subtract the latter set of difference scores from the former to obtain an estimate of viewpoint dependent changes in the balance of response strategies (i.e. any shift to a greater reliance on mental rotation). As readers can see, there is no evidence for a robust relationship between any such shift and the typical vividness of people’s visualisations (r = 0.11, p = 0.33; see Figure 6a).

**Figure 6.**
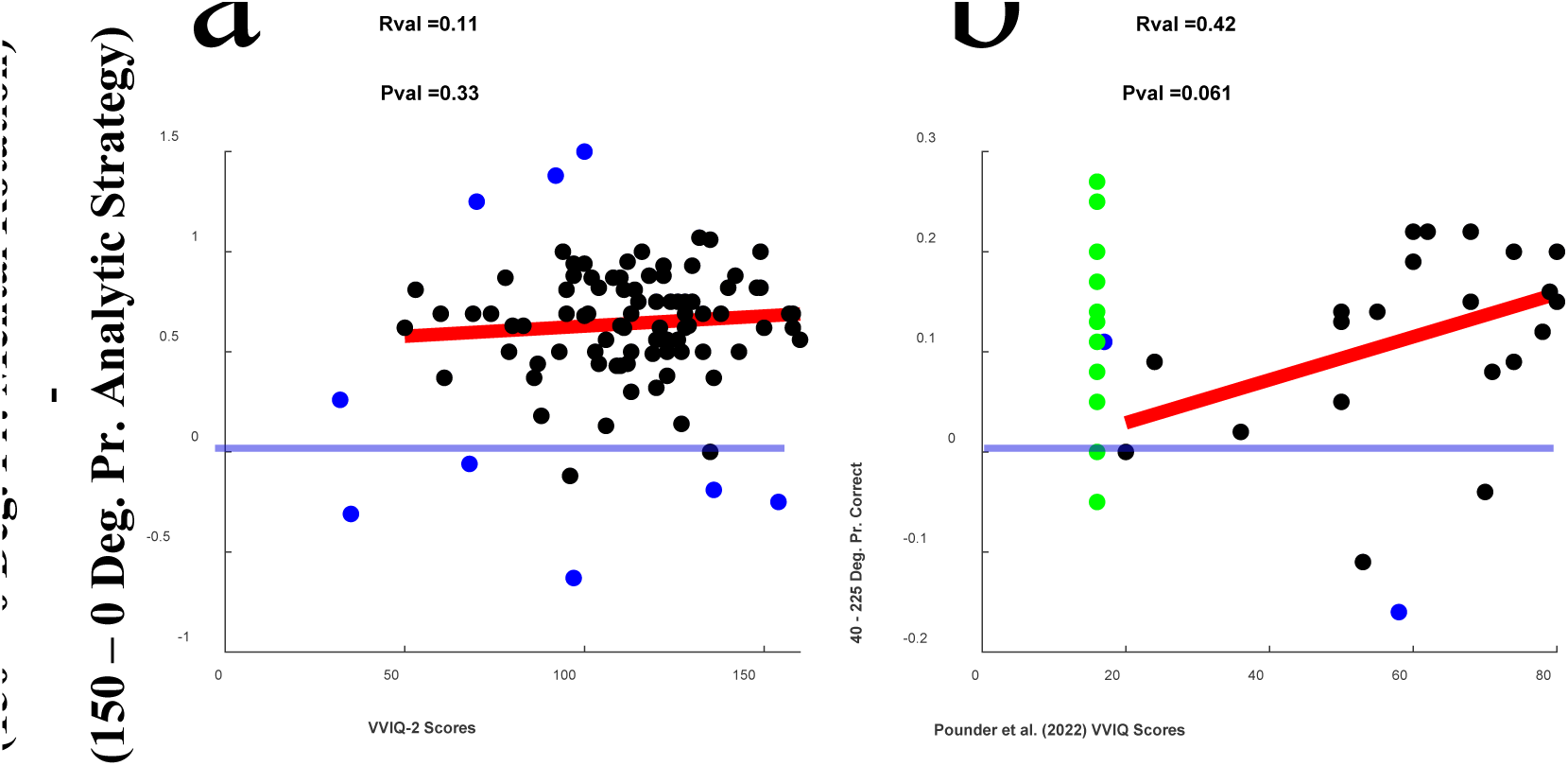
**a)** Viewpoint dependent changes in response strategy. We calculate difference scores relating to the probabilities of reporting mental rotation when objects are depicted from different, as opposed to the same viewpoint. We also calculate difference scores relating to probabilities of reporting an analytic response strategy, when objects are depicted from different as opposed to the same viewpoint. We subtract the latter set of difference scores from the former (Y-axis). Individual VVIQ-2 scores are plotted on the X-axis. **(b)** Data re-plotted and analysed from Pounder et al. (2022). On the Y-axis we plot differences in the proportion of correct decisions when objects were depicted from viewpoints horizontally rotated by 225° as opposed to 40°. Aphant datapoints (people achieving the minimum possible VVIQ-2 score) are coloured green, and are excluded from analyses. Remaining data are treated in the same way as data in all our correlational analyses, and are most comparable to data depicted in Figure 3a.

At first, this last finding might seem counter intuitive. The confusion is resolved by understanding that the typical vividness of people’s visualisations also predicted what response strategy they would use on trials when objects were depicted from the *same* viewpoint (Pr. Mental Rotation x VVIQ-2 scores, r = 0.25, p = 0.02). So, there is no robust increase in the probability that people with vivid imagery would rely on mental rotation, as they were already likely to be using mental rotation on baseline trials.

### Descriptions of strategies that participants felt had not been available as a response option

Fifteen participants included in data analyses took the opportunity to report using a response strategy that we had not described. Of these descriptions, we regarded seven as having described another form of visualisation. Three of these involved imagining viewing objects from different viewpoints (e.g. “*I imagined myself standing looking at each object from a certain perspective and comparing them*”). One described a more complex form of visualisation (“*Following one side of the shape from one end to other. Do so to both and compare visualised line*”), and one described a form of motor imagery (“*Careful examination of the 3D shape, alongside hand gestures to see the direction of the cubes*”). The other two were differently worded descriptions of visualisations (e.g. “*I rotated it around*”).

We regarded another seven responses as having described a different analytic strategy. Three highlighted symmetry detection (e.g. “seeing they were a reflection of each other, which means they cannot be rotated to appear the same”), three described a counting strategy (e.g. “*Count blocks going in each direction*”), and one described an analytic strategy involving the comparison of prominent features (“…*if they are have a bend on the same side when they are 180 different then they are different*”). The final description described confusion (“*The image on the right is not fully displayed, so I can’t confirm if they are the same*”). These data highlight that people adopt a range of visualising and analytic response strategies when completing a mental rotation task.

## Discussion

Our data suggest that viewpoint dependent changes in performance on mental rotation tasks are only *weakly* associated with the subjective vividness of people’s visualisations (see Figure 3), or with reliance on mental rotation as a response strategy (see Figure 4). Mental rotation was a superior response strategy when judging if objects viewed from different viewpoints were the same or different (see Figure 4b), but the vividness of people’s visualisations (VVIQ2 scores) did not predict if they would benefit from this strategy (see Figure 6a). People with vivid imaginations were likely to report relying on mental rotation on baseline trials involving different, mirror reversed objects – limiting their capacity to benefit from an increased reliance on this strategy when objects were depicted from different viewpoints. Decision response times were not associated with the vividness of people’s visualisations (see Figure 3a and 5) or with reliance on mental rotation as a response strategy (see Figure 4a).

### Why did people report visualising when objects were depicted from the same viewpoint?

Many of our participants (93%) reported visualising an object rotating on a subset of trials (M 25%, SD 14) in which *different* objects had been depicted from the same viewpoint. We can be confident these visualisations were selective to trials depicting *different* objects due to a feature of our study design, and because of how we have treated data.

Our task involved a 2-stage response cycle, with people first reporting on if they had visualised an object rotating. They were only asked for a second response if they did not endorse this first option. This encouraged participants who were primarily motivated to rapidly finish the study to endorse the first option – even when that did not make sense (i.e. when test images showed the *same* object depicted from the *same* viewpoint, see Figure 1a). We therefore removed *all* participants from analyses who had indicated that they had visualised an object rotating on any of these trials. Note that this will not only have removed people who were primarily motivated to rapidly finish the study, it will also have removed some people who were inattentive. So, by definition, any remaining reports of visualising an object rotating when test images depicted objects from the same viewpoint were related to a presentation of *different* mirror reversed objects (e.g. see Figure 1b). We are confident these responses were systematic, as opposed to an inattentive issue, as the remaining participants, who had *never* reported visualising when matching objects had been depicted from the same viewpoint, *often* reported visualising when different objects had been presented in the same circumstances (M = 25%, SD = 14, range = 0 – 50%). We regard this as strong evidence that, in the context of a mental rotation task, people will often attempt to use visualisation to decide if *different* mirror reversed objects are matched, even if the different objects are depicted from the same viewpoint (see Figure 1a).

Other researchers have been mindful of this possibility, so they have adopted the practise of excluding trials that depict different (mirror reversed) objects from analyses that estimate the speed of mental rotations (e.g. Hilton et al., 2022; Searle & Hamm, 2017). We did not exclude such trials because a key point of interest was the propensity of people to use different response strategies.

A clear implication to other researchers is, however, that trials that involve different objects depicted from the same viewpoint will likely involve some visualisations. These data have often been treated as a visualisation free ‘baseline’, that can either be subtracted from data from all other conditions (e.g. Zhao et al., 2022), or which can be used as an anchor point, theoretically devoid of the time required to visualise an object rotating, from which a slope involving datapoints from other conditions can be calculated to estimate mental rotation speeds (e.g. Gardony et al., 2014; Kail & Park, 1990; Shepard & Metzler, 2971; Zeman et al., 2010). In either case, the inadvertent inclusion of visualisations within a putative baseline could distort estimates of the propensity to visualise, and impact of visualisations.

### Mental rotation and Aphantasia

The data we have reported on only involved one person whom we would regard as an Aphant (i.e. a person who obtains a minimum score of 32 on the VVIQ-2 – showing that they rated each experience as ‘*No image at all, you only “know” that you are thinking of the object*’).

There were only six people who rated their imagined experiences as vague and dim, or as ‘*No image at all…*’ which corresponds with ∼6% of our sample – consistent with estimates of the prevalence of vague and dim imagery across the general population (Dance et al., 2022). Nonetheless, we can ask how our data relate to studies that have targeted Aphants?

Our data are broadly consistent with some aspects of a recent study, that tested for differences on mental rotation between self-reported Aphants and controls (Kay et al., 2024). While Aphants were overall *more* accurate, they were also slower to make decisions. These data are admittedly ambiguous, as they are consistent with a common strategy but a more cautious approach within the confines of a speed accuracy trade-off (Heitz, 2014).

Nonetheless, Aphants were more likely to report using an analytic response strategy, and that is certainly consistent with the people who tended to report experiencing low vividness imagery in our sample.

Another study found little evidence for a difference in mental rotation decision accuracy between Aphants and a control group, but Aphants tended to be slower to make decisions (Pounder et al., 2022). So, there is some consistency between the two studies that have targeted groups of Aphants in this context (Pounder et al., 2022; Dance et al., 2022). We also noticed what we thought was an interesting parallel in data reported by these researchers (Pounder et al., 2022) and our own data. We have re-plotted a subset of their mental rotation data in Figure 6b (we thank the authors for making these data available), adjusted to match the format of our own data as closely as possible. To this end, we calculated proportion correct difference scores between the largest (225°) and smallest (40°) viewpoint differences they reported on, and we correlate these with VVIQ scores. We colour Aphant data points green and exclude these from analyses. We then treat their remaining data as we have our own, and find a trend matching our result – with people who reported more vivid imagery tending to make proportionally fewer incorrect decisions about objects depicted from more different viewpoints. Admittedly this trend is not robust (p = 0.061), but it is stronger than our effects (r = 0.42, as opposed to 0.3) and so it is plausible that the different outcomes reflect on the different sample sizes (23 in our re-analysis of their data, 86 in the corresponding analysis of our own data).

We have laboured this last point because it is possible that our sample of the general population has reached a similar outcome as the prior study (Pounder et al., 2022), provided that people who were clearly Aphants are excluded from analysis. The implication would be that Aphants may be separable from the general population, in terms of how the subjective vividness of their imagery interacts with mental rotation task performance.

### Do people with Aphantasia lack insight into their capacity to visualise?

It has been suggested that people with Aphantasia might be able to have imagined visual sensations, which guides their performance on mental imagery tasks, but lack insight into this capacity (Nanay, 2021; Siena & Simons, 2024; Pounder et al., 2022). We think this is unlikely.

As self-described Aphants (Bouyer & Arnold, 2024), Arnold and Bouyer have very different subjective experiences of mental rotation tasks. However, in each case this seems to involve an easily recognised and reportable form of mental exertion. Arnold resorts to a time-consuming cross examination of the paired images, searching for a turning point that is logically inconsistent across the two images (i.e. where one has a right turn, the other a left). To him, mental rotation tasks are a time-consuming logical puzzle. Bouyers’s experiences can be described as involving imagined texture and motor sensations. She imagines how an object would feel in her hand, and the sensation of adjusting it (similar to the description of one of our participants). She feels that this gives her insight into whether the two objects are likely the same or different. Bouyer’s experiences clearly involve imagined sensations, and they might tap common processes relative to those that promote visual imagery in other people. She does, however, have a sense of insight into her imagined sensations, and can describe them – and they are not at all visual in nature. This highlights an established theme of heterogeneity in relation to the experiences of Aphants, and cautions against assuming they will all perform similarly on tasks that attempt to measure imagery (also see Dawes et al., 2020; Takahashi et al., 2023).

We recommend that researchers ask about the strategies people use when performing imagery related tasks, and predict that diverse and variably effective strategies will be reported by Aphants and non-aphants. We join others in arguing that at this early stage of Aphantasia research, it is important to gather qualitative information about how people experience their thoughts, in the hope that this will inform more objective tests and investigations (also see Dawes et al., 2020; Dawes et al., 2024).

## Conclusions

Our data suggest that viewpoint dependent changes in performance on mental rotation tasks are only *weakly* associated with the subjective vividness of people’s visualisations, or with reliance on mental rotation as a response strategy. While mental rotation was a superior response strategy, the vividness of people’s visualisations did not predict if they would benefit from that strategy. We suggest that better metrics of visualisation are needed, and suggest it is important to enquire about the response strategies people use when performing imagery related tasks.

## Acknowledgements

This research was supported by a Discovery Project Grant, funded by the Australian Research Council, awarded to D.H.A.

